# Slow-wave sleep is associated with nucleus accumbens volume in elderly adults

**DOI:** 10.1101/2024.10.25.620231

**Authors:** Kitti Bán, Ádám Nárai, Noémi Báthori, Éva M. Bankó, Adél Bihari, Vivien Tomacsek, Tibor Kovács, Béla Weiss, Petra Hermann, Péter Simor, Zoltán Vidnyánszky

## Abstract

Slow-wave sleep (SWS) is essential for restorative neural processes, and its decline is associated with both healthy and pathological ageing. Building on previous rodent research, this longitudinal study identified a significant association between nucleus accumbens (NAcc) volume and SWS duration in cognitively unimpaired older adults. Our findings support the involvement of the NAcc in ageing-related modulation of SWS and suggest potential therapeutic targets for improving SWS.

## Introduction

Proper regulation of sleep and wakefulness is essential for maintaining overall health, cognitive function, and emotional well-being^1^. With age, overall sleep duration, efficiency, rapid eye-movement (REM) sleep, and non-rapid eye-movement (NREM) sleep are reduced whilst sleep-wake fragmentation increases, which disruptions are further amplified in neurodegenerative disorders (NDs)^2,3^. Decreased slow-wave sleep (SWS), also known as deep NREM sleep or N3 sleep, plays a pivotal role in fundamental neural mechanisms; it is associated with restorative processes, including cellular maintenance^4^, glymphatic clearance^5,6^, renormalisation of synaptic weights^7^, and memory consolidation^8,9^.

Human neuroimaging studies investigating the relationship between brain structure and SWS macroarchitecture (duration, percentage of sleep time) revealed supporting evidence for the link between reduced SWS and markers of accelerated brain-ageing and increased neurodegenerative risk, including decreased cortical^10,11,12^ and subcortical^10,13^ volume as well as increased white matter hyperintensities^10^. Indeed, SWS and brain atrophy are suggested to share a bidirectional relationship, whereby brain atrophy reduces SWS, which disrupts regenerative neural processes, in turn leading to more atrophy^14,15^.

Recent research in mice have identified the nucleus accumbens (NAcc), a brain structure critical for motivation, emotion, learning, and motor function, as a central neuromodulatory hub involved in the regulation of slow-wave sleep (SWS)^16,17.18.19^. However, evidence supporting this link in humans remains lacking. To address this gap, we conducted a comprehensive longitudinal investigation of the association between NAcc morphology and SWS in a healthy ageing population. To this end, we utilised multi-night, at-home electroencephalogram (EEG) recordings to examine SWS macroarchitecture and structural magnetic resonance imaging (MRI) to measure NAcc morphology (Fig.1). By examining the relationship between these measures, this study aimed to elucidate the potential role of the NAcc in the regulation of SWS in humans and its implications for healthy ageing.

**Figure 1.**
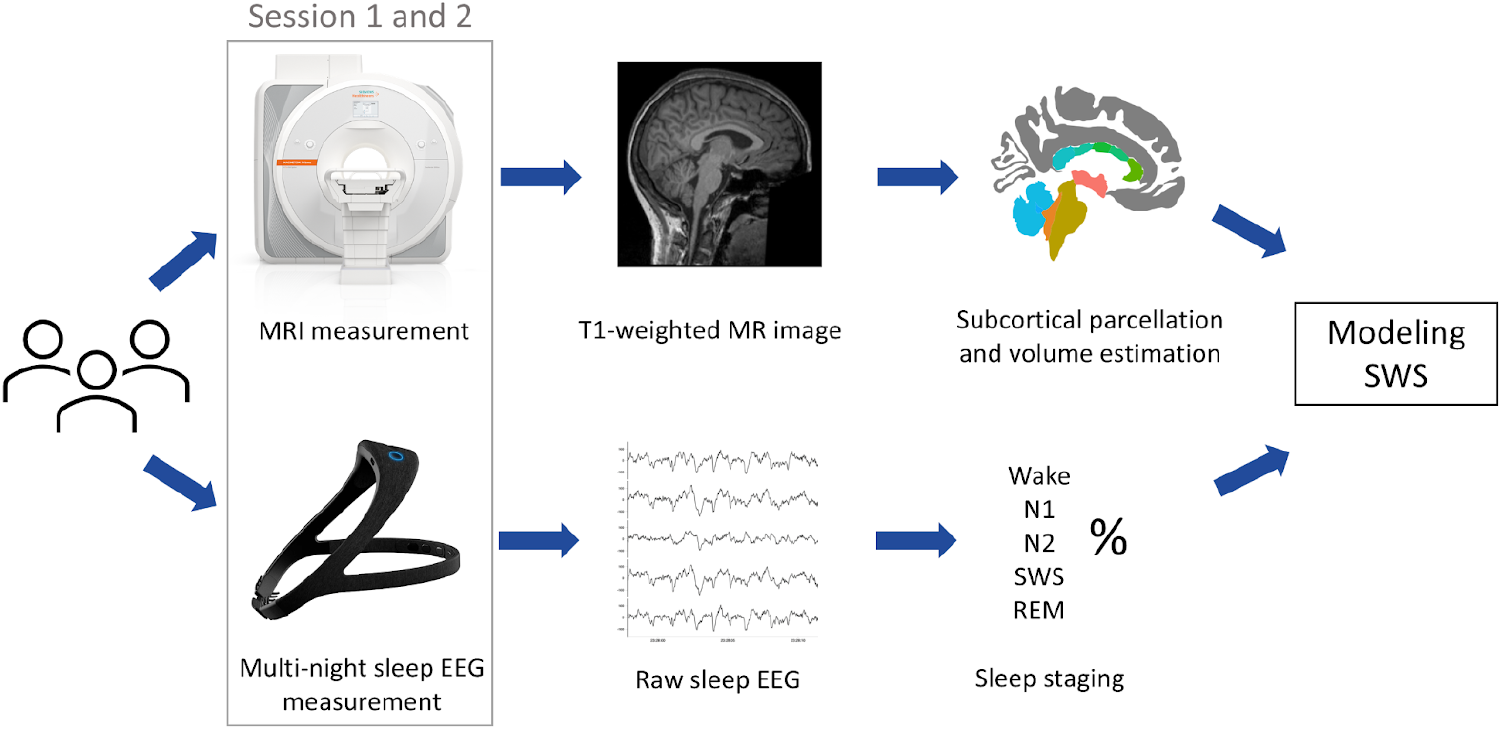
Schematic representation of data collection and analysis protocol. Participants between the ages of 50-80 years underwent structural MRI scanning and EEG sleep recording using the Dreem2 headband for a 7-day period at home in two sessions 1.5 years apart (*N* = 64 in Session 1, *N* = 48 in Session 2, *N* = 43 in both sessions). High-resolution 3D T1-weighted structural images were used to extract the volume of 8 subcortical regions. Dreem’s machine-learning algorithm divided sleep data into the following stages: wake, N1 sleep, N2 sleep, N3 sleep, SWS, and REM sleep. For each participant, each sleep stage duration was averaged across all measurement days per session. We utilised linear modelling to investigate the association between SWS percentage and the volume of the 8 subcortical areas whilst controlling for age, sex, session, motion-related Euler number, and intracranial volume.

## Results

Linear mixed effect modelling using all participants’ data from both sessions (*N* = 112) revealed a significant association between SWS percentage and NAcc volume (*F* = 4.27, *p* = .041), session *(F* = 4.56, *p* = .037), age (*F* = .077, *p* = .78), sex (*F* = 3.27, *p* = .075), ICV (*F* = .34, *p* = .56), and the motion-related Euler number (*F* = .61, *p* = .44). The significant link between SWS percentage and NAcc volume was confirmed (*F* = 4.77, *p* = .032) using the same model and covariates (for all, *p* > .05) but only including data from the subset of participants with valid measurements from both sessions (*N* = 86).

Given that session type showed significantly higher percentage of SWS in Session 1 when including all participants, we also examined the association between SWS percentage and NAcc volume separately for each session. Linear modelling with the same covariates as above revealed a significant link between SWS percentage and NAcc volume for both Session 1 (*N* = 64, *F* = 10.41, *p* = .0026) and Session 2 (*N* = 48, *F* = 9.48, *p* = .0039). The effects of covariates were not significant (*p* > .05) in either of the models, except for sex in Session 2 (*F* = 202.02, *p* = .032). The consistent relationship between NAcc volume and SWS percentage across sessions is further underlined by the strong Spearman correlation between SWS percentage (*N* = 43, *R*_*s*_ = 0.78, *p* < .0001) from sessions 1 and 2 as well as NAcc volume (*N* = 43, *R*_*s*_ = .84, *p* < .0001) from sessions 1 and 2 (Fig. 2b).

**Figure 2.**
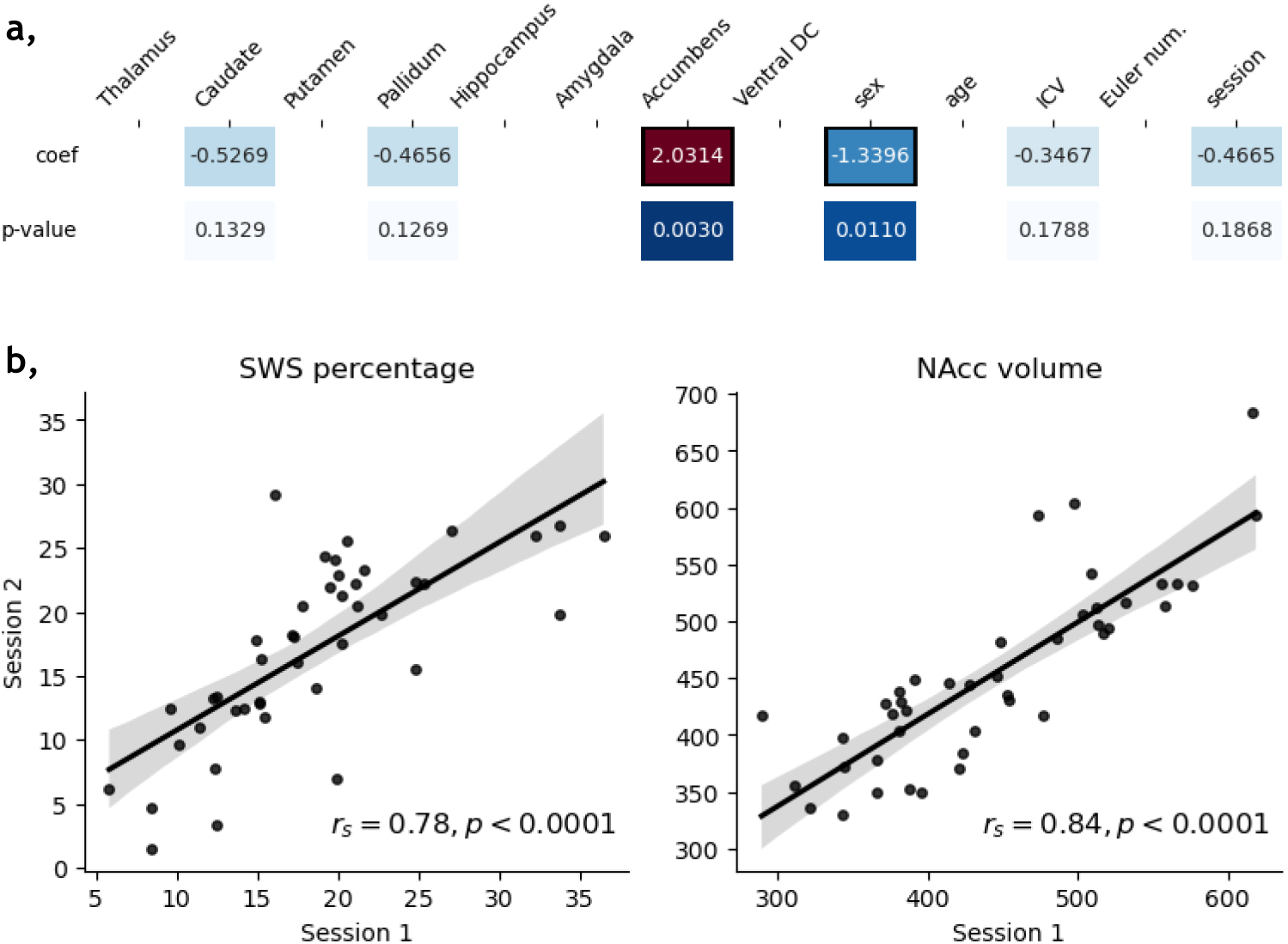
Analyses of SWS percentage and NAcc volume. **a**, Investigation of the relationship between 8 subcortical regions and SWS percentage whilst controlling for sex, age, intracranial volume (ICV), Euler number, and session type using an Elastic-Net regularised linear model. White cells indicate zero coefficients and cells with black frames represent a significant association (*p* < .05) with SWS percentage. **b**, The strong correlation between SWS percentage across sessions (left) and NAcc volume across sessions (right) demonstrates the robustness of these measures. Each graph shows the least-squares fit line and its 95% confidence band as the shaded area. The Spearman correlation coefficient (*r*_*s*_) and its corresponding *p*-value (*p*) are shown on the bottom right of each graph.

Our extended linear model with Elastic-Net regularisation (*N* = 112) investigated the relationship between all 8 subcortical regions and SWS percentage whilst controlling for the same covariates as our previous models (Fig.2a). Only NAcc volume (β = 2.03, *p* = .003) and sex (β = −1.34, *p* = .011) showed a significant association with SWS percentage. Thalamus, caudate, putamen, pallidum, hippocampus, amygdala, ventral diencephalon (DC) volumes, as well as age, ICV, Euler number, and session were not significantly (*p* > .05) related to SWS percentage.

## Discussion

This longitudinal study showed a significant positive association between NAcc volume and SWS percentage, which link was robust across measurement sessions. This result is consistent with evidence from rodent models that found the NAcc to regulate SWS^16,17.18.19^. Similarly, existing human neuroimaging studies indicate a significant correlation between SWS and subcortical areas, including overall subcortical volume^10^ and the thalamus^13^. To our knowledge, only one study^20^ examined the relationship between SWS and NAcc volume, which reported a lack of significant connection between the two measures. The discrepancy between these and our results likely stems from methodological differences. Specifically, Weihs and colleagues used a different MRI scanning protocol with a lower magnetic field strength, examined the link between SWS and brain volumes cross-sectionally, and utilised single-night, lab-based polysomnography (PSG) to quantify SWS.

The precise ways through which the NAcc regulates sleep in humans is unclear, although rodent studies are able to shed some light on potential cellular mechanisms. Selective activation of A_2A_ adenosine-expressing^18,19^ and D_2_ dopamine-expressing^16,17^ neurons in the NAcc were found to strongly promote SWS in mice, whilst D_1_-expressing neurons induced behavioural arousal^17^ and REM sleep^16^. As D_2_ and A_2A_ receptors are co-expressed in striatopallidal neurons, it is hypothesised that the NAcc modulates sleep and wakefulness through dopaminergic input from the VTA, which conveys arousal-promoting motivational information^19^. Together with rodent studies, our results further imply the NAcc as a pivot point for balancing sleep and wakefulness through complex neuromodulatory circuits throughout the midbrain, basal ganglia, and cortex. Future studies are necessary to confirm how the NAcc may reciprocally regulate sleep and motivational behaviour in humans.

Our results have important implications for understanding the neural basis of SWS deterioration in ageing. Ageing is associated with a decline of the brain’s dopaminergic system^21^, an increased risk of NDs^22^, and heightened levels of chronic stress^23^, which factors exacerbate one another^15,23,24^. The NAcc is a key hub integrating motivational, sensorimotor, affective, and cognitive information within the dopaminergic network^25^ and is particularly vulnerable to atrophy in NDs and chronic stress^26^, both of which are linked to sleep disturbances^2,16^. Our findings suggest that age-related deterioration of the NAcc may play a critical role in SWS impairment in older adults and raise the prospect that NAcc structural and functional integrity might serve as a neural marker or therapeutic target for ageing-related SWS decline.

## Methods

### Participants

Community-dwelling older middle-aged and older adults between the ages of 50 and 80 years were recruited as part of the Hungarian Longitudinal Study of Healthy Brain Aging (HuBA) carried out at the Brain Imaging Centre of the HUN-REN Research Centre for Natural Sciences in Budapest, Hungary. All participants provided written informed consent prior to the study, which was approved by the National Institute of Pharmacy and Nutrition (OGYÉI/68903/2020) and was conducted in accordance with the Declaration of Helsinki. The study protocol, exclusion and inclusion criteria, and participant demographic details are described in this paper^27^.

Participants underwent the same measurement protocol two times (Session 1 and 2 from now on), 1.5 years apart. In this study, we used data from participants who had both structural MRI scans and sleep recordings in at least one of the two sessions. Session 1 data from five participants and Session 2 data from two participants were excluded due to having less than 2 nights of sleep recordings of sufficient quality. The final sample included 64 participants (33 females, *M*_*age*_ = 62.2 years, *SD*_*age*_ = 6.9 years) in Session 1 and 48 participants (27 females, *M*_*age*_ = 63.7 years, *SD*_*age*_ = 6.6 years) in Session 2. A total of 43 participants had measurements from both sessions (23 females, *M*_*age*_ = 61.9 years, *SD*_*age*_ = 6.7 years in Session 1).

### Sleep data acquisition

The wireless Dreem2 headband (DH; Rhythm, Paris, France) was used to collect electrophysiological data during participants’ sleep at home for a 7-day period. This device recorded electric signals from the scalp at a sampling frequency of 250 Hz utilising 5 dry electrodes and a frontal ground electrode (Fp2), resulting in seven bipolar derivations; F7-O1, F8-O2, Fp1-F8, F8-F7, Fp1-O1, Fp1-O2 and Fp1-F7. The headband also incorporated a 3D accelerometer measuring body movements and an infrared pulse sensor, both sampling at 50 Hz. We selected the DH to measure sleep macroarchitecture as it has been indicated as a reliable and practical device to record sleep EEG for multiple nights in the home environment^28,29,30^. Importantly, data provided by the DH was found comparable to those obtained with traditional PSG recordings^31^.

During one-to-one lab visits, participants received the DH together with detailed instructions on how to use the device and its accompanying mobile application. Participants were also walked through a short sample recording session, during which they were taught how to properly position and charge the device, start and stop recordings, and upload the data to Dreem’s servers. To provide off-site assistance, participants received a detailed, paper-based manual as well as instruction videos on how to correctly use the device. To enhance signal quality, participants were equipped with isopropyl alcohol wipes to clean their skin and sensors on a daily basis as well as a sweatband designed to secure the DH closer to the skin and the scalp.

During the 7-day measurement period, participants were disclosed no information about the recordings and in case additional support was required, they were guided through emails or video calls to optimise headband placement and recording quality. During a second lab visit, participants returned the headband and had the opportunity to ask for a brief summary of the study and view the hypnograms illustrating the quality of their sleep. The equipment and procedures used were identical across the two data collection sessions.

### Sleep data preprocessing

In addition to recording physiological signals, the DH offers an automated, machine-learning-based sleep staging algorithm, which qualifies and scores sleep recordings. As noise and artefacts may affect sleep staging, the algorithm also assigns a quality index to each EEG channel, which represents the corresponding signal-to-noise ratio. This automatic analysis ensures the consistency of sleep staging across epochs and recordings, whereas manual scoring can greatly vary among sleep experts, introducing large inter-rater variability^32^. Sleep scoring by Dreem’s algorithm was demonstrated to show reliable performance akin to human expert scoring of PSG data^31^.

To ensure the reliability of this automated sleep staging, a qualified PSG expert (P.S.) visually assessed 20 randomly selected whole-night (>5 hours) recordings using visual EEG scoring rules^33^. This procedure allowed for the establishment of sleep hypnograms, comparison between the manually- and automatically-derived scores, and identification of a quality threshold for the study. The Pearson correlation between automatic and manual scorings ranged between .79 – .90 (Wake: .64 – .85, N1 sleep: .11 – .77, N2 sleep: .81 –.96, SWS: .59 – .94, REM: .67 – .98) in cases when signal quality was at least 75% in at least one channel but dropped below .60 if signal quality was poorer. Consequently, we constrained further analyses to recordings in which signal quality exceeded a threshold of 75% in at least one channel. After averaging each participant’s available sleep measures over all nights in each session, we used mean SWS percentage for further analyses.

### MRI acquisition and preprocessing

High-resolution 3D T1-weighted structural images were acquired on a Siemens Magnetom Prisma 3T MRI scanner with a 32-channel head coil using the following parameters: isotropic 1 mm^3^ spatial resolution, field of view (FOV) = 256 × 256 mm, 2-fold in-plane GRAPPA acceleration, repetition time (TR) = 2300 ms, echo time (TE) = 3 ms, inversion time (TI) = 900 ms and flip angle (FA) = 9°.

Subcortical segmentation of T1-weighted structural images and estimation of morphometric statistics were performed using FreeSurfer 7.1.1 with the built-in probabilistic atlas^34^. For our analyses, we extracted the volumes of eight subcortical ROIs (Thalamus, Caudate, Putamen, Pallidum, Hippocampus, Amygdala, Accumbens, and Ventral Diencephalon) averaged across hemispheres and intracranial volume (ICV).

### Statistical analyses

We utilised linear models to uncover the link between NAcc volume and SWS percentage whilst controlling for age, sex, ICV, and Euler number (calculated from the FreeSurfer output). As the Euler number has been shown to be highly correlated with manual ratings of motion artefacts^35^, it was included in our analyses to account for the possible effect of participant motion.

A linear mixed effects (LME) model including measurement data from both sessions was used to investigate the association between SWS percentage and NAcc volume. Fixed effects were added for NAcc volume, age, sex, ICV, Euler number, and session, whilst a random intercept for participants was also included;

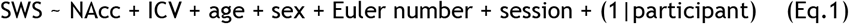

As this model showed a significant effect of session type on SWS percentage, we implemented separate linear models for Session 1 and Session 2 data to confirm the link between SWS percentage and NAcc volume. Similarly to the first model, these analyses also included covariates for age, sex, ICV, and Euler number;

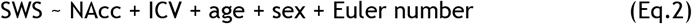

The above models were implemented in R (v. 4.3.3)^36^ via the lme4 (v. 1.1.35.3)^37^ and stats (v. 4.3.3)^36^ packages.

Our final analysis examined whether SWS percentage is uniquely linked to the NAcc or was also associated with the volume of other subcortical regions, including the thalamus, caudate, putamen, hippocampus, amygdala, or ventral diencephalon (DC).

To account for the relatively high number of predictor variables (8 subcortical regions, age, sex, ICV, and Euler number) in this analysis, we implemented linear regression models with Elastic-Net regularisation, with both L1 and L2 penalty terms added. Hyperparameter tuning was performed via 5-fold cross-validation with 20 *l1_ratio* values between 0 and 1 on an inverse logarithmic scale and 20 automatically selected values for regularisation strength. Models were refitted and evaluated on the whole dataset to calculate the final coefficients. Statistical significance of coefficients was determined using two-tailed permutation tests with 1000 random permutations and alpha level of .05. All features were standardised to aid the interpretability of coefficients. Modelling was performed in Python (v. 3.10.13) using the scikit-learn package (v. 1.4.1)^38^.

## Acknowledgements

This research was supported by the National Brain Research Program 3.0. by the Hungarian Academy of Sciences (NAP2022-I-1/2022; PI: Z.V.) and the Hungarian Research Network (HUN-REN; 298/4/2023/HF; PI: Z.V.). P.S. was supported by the Hungarian National Research, Development, and Innovation Office Grant (NKFI FK 142945) and by the Janos Bolyai scholarship of the Hungarian Academy of Sciences.

B.W. was supported by the Janos Bolyai scholarship of the Hungarian Academy of Sciences. We thank Emília Nagy, Annamária Manga, István Hevesi, Menta Havadi-Nagy, Rebeka Kelemen, and Ádám Simon for taking part in data collection.

## Author contributions

Z.V. acquired funding and conceptualised the study. Z.V., P.H., Á.N., P.S., N.B., and T.K. designed the study. N.B., V.T., and A.B. collected the data. Á.N., P.H., P.S., N.B., É.B., B.W., and K.B. analysed the data. Á.N. prepared the figures. K.B., Á.N., and Z.V. wrote the manuscript. P.H., P.S., N.B., V.T., É.B., A.B., B.W., and T.K. edited the manuscript.

